# Cyclophilin A is a mitochondrial factor that forms antiapoptotic complexes with p23

**DOI:** 10.1101/2020.08.21.261982

**Authors:** Cristina Daneri-Becerra, Brenda Valeiras, Mariana Lagadari, Mario D. Galigniana

## Abstract

Cyclophilin A (CyPA) is an abundant and ubiquitously expressed protein belonging to the immunophilin family that has intrinsic peptidyl-prolyl-(*cis/trans*)-isomerase enzymatic activity. In addition to mediating the immunosuppressive effects of the drug cyclosporine A, CyPA is involved in multiple cellular processes such as protein folding, intracellular trafficking, signal transduction, and transcriptional regulation. Because CyPA is also a molecular chaperone, its expression is induced by several stressor agents and is a highly abundant protein in cancer cells. In this study, it is demonstrated that in several cell types and at least in murine liver, a significant pool of this immunophilin is primarily an intramitochondrial factor that migrates to the nucleus upon the onset of stress. It is also shown that CyPA has antiapoptotic action. Importantly, the capability of CyPA to form complexes with the small acidic cochaperone p23 is proven, this interaction being independent of the usual association of p23 with the heat-shock protein of 90-kDa, Hsp90. Furthermore, it is demonstrated that the CyPA•p23 complex enhances the antiapoptotic response of the cell, suggesting that both proteins form a functional unit whose high level of expression plays a significant role in cell survival.

## Introduction

Cyclophilin A (CyPA) is an abundant protein belonging to the immunophilin (IMM) family. As such, it is characterized by the presence of a signature domain, the peptidyl-prolyl isomerase (PPIase) domain, that shows the capability to catalyze *cis*↔*trans* isomerization of peptidyl-prolyl bonds (Nigro et al., 2013; Zgajnar et al., 2019). CyPA was isolated from bovine thymocytes and identified as the cytosolic receptor for the immunosuppressive cyclic undecapeptide cyclosporin A, such that the formation of CyPA•drug complex impairs the transcription of genes related to the immune response and prevents proliferation of T-cells (Handschumacher et al., 1984).

CyPA is ubiquitously expressed in the cell, and can also be secreted to the medium in response to inflammatory stimuli such as hypoxia, infection, and oxidative stress, among others (Nigro et al., 2013). Importantly, the secreted form shows autocrine and paracrine properties by binding to the CD147 cell surface receptor (Pushkarsky et al., 2001). CyPA shows pleiotropic actions that comprise protein folding, trafficking, cell activation, cardiovascular diseases, chemiotaxis, atherosclerosis, diabetes, viral infections, drug resistance, cancer, neurodegenerative diseases, ageing, etc. (Dawar et al., 2017; de Wilde et al., 2018; Nigro et al., 2013; Xue et al., 2018). Interestingly, CyPA binding to CD147 has been related to prosurvival pathways in neuronal cultures (Boulos et al., 2007; Yurchenko et al., 2002).

Even though CyPA may play relevant roles during protein folding and the induction of conformational changes of various proteins thanks to its PPIase activity, it also functions as a standard molecular chaperone modulating protein-protein interactions, regardless of its intrinsic enzymatic activity (Rein, 2020). Thus, CyPA has been related to the active cytosolic mechanism of retrotransport of soluble proteins via dynein in a PPIase domain-dependent and enzymatic activity-independent manner (Galigniana et al., 2004). Several studies have also demonstrated that CyPA induces leukocyte chemotaxis through direct binding to the ectodomain of receptor CD147 in a PPIase-independent fashion (Song et al., 2011). Similarly, CyPA impairs the early stage of influenza A virus replication by simple protein-protein interaction promoting the degradation of the virus matrix protein M1 (Mahesutihan et al., 2018).

Inasmuch as many PPIases are also molecular chaperones, it has recently been proposed that the association of this family of proteins to client-factors can influence the energy landscape by destabilization of intermediates and unfolding activity (Galigniana, 2020; Rein, 2020). These events may be totally independent of the enzymatic activity depending on the specific client-protein. In this regard, it should be pointed out that molecular chaperones often act in a sequential manner. For example, chaperones associated to steroid receptors and protein kinases are usually transferred from the Hsp70 folding machinery to the Hsp90-based folding platform (Pratt et al., 2004b), which replaces intermediate complexes containing the co-chaperone Hop/p60 by a high molecular weight IMM. The entire heterocomplex is finally stabilized by the small acidic Hsp90-binding co-chaperone, p23 (Pratt et al., 2004a). It is accepted that the overexpression of Hsp70 or Hsp90 inhibits apoptosis and prevents caspase activation in many cellular models upon accumulation of misfolded proteins, oxidative stress or DNA damage (Lanneau et al., 2008; Mosser and Morimoto, 2004). The recruitment or overexpression of the co-chaperone p23 should consequently contribute to this outcome. Accordingly, it has been recently reported that both apoptosis and cell proliferation are augmented in fibroblasts where the expression of p23 was knocked-out (Madon-Simon et al., 2017). Because the phenotype of p23-null embryos is like the one observed when the glucocorticoid receptor function is abolished, it was speculated that the reason is based on the impact p23 on this particular transcription factor.

CyPA has also been related to the mechanism of apoptosis by favouring the nuclear translocation of the Apoptosis-Inducing Factor (AIF) (Zhu et al., 2007). In vitro, recombinant AIF and CyPA cooperate in the degradation of plasmid DNA and induce DNA loss in purified nuclei. However, CyPA down-regulation does not arrest nuclear translocation of AIF, but reduces DNA damage (Artus et al., 2010). In an alternative model, it was shown that one of the first responses to oxidative stress involves the rapid accumulation of CypA in the nucleus some hours before apoptosis is triggered (Doti et al., 2014). Targeting of CypA with synthetic peptides to inhibit the AIF-mediated neuronal loss was also used as a valuable strategy to protect cells (Rodriguez et al., 2020). This raises the possibility that endogenous factors capable to interact with CyPA may also function as protective factors against apoptosis, a seductive hypothesis that is examined in this study.

On the other hand, CyPA has been shown to be implicated in essential neuronal functions such as axonal transport, synaptic vesicle assembly, neuroprotective roles against abnormal protein aggregation (Avramut and Achim, 2003), and as a necessary factor related to the BDNF neuroprotective action (Jeon et al., 2008; Lin et al., 2014). Also, it has been suggested that CyPA may play a neuroprotective role against both oxidative stress (Doyle et al., 1999) and ischemic events (Boulos et al., 2007). Moreover, it has also been reported that the direct injection of purified CyPA exerts protective roles after brain injury (Redell et al., 2007). In other words, CyPA influences a wide variety of biochemical and pathophysiological pathways in several models of disease, often in opposite manner, its effects being related to the type of complexes CyPA participates.

In preliminary confocal microscopy studies where fresh cells were observed, we noted that CyPA shows a subcellular fluorescent pattern quite compatible with a primary mitochondrial localization. During these observations, it was evidenced a strong colocalization of CyPA with the Hsp90-binding co-chaperone, p23. In view of the above-referenced reports showing that both proteins may individually participate in apoptotic pathways, we hypothesize that it could be entirely possible that CyPA is a mitochondrial factor that impacts on apoptosis itself, and that both chaperones CyPA and p23 may belong to heterocomplexes that could influence cell survival. In this work it is demonstrated that CyPA is a novel mitochondrial factor with antiapoptotic properties, which are enhanced by the interaction of CyPA with the cochaperone p23 in an Hsp90-independent manner.

## Results

### CyPA is a mitochondrial factor

Confocal microscopy images from several cell types showed that CyPA (green) exhibits a profile compatible with mitochondrial localization (Fig.1-A). Accordingly, colocalization of this IMM and the specific mitochondrial fluorescent probe MitoTracker (red) was also evidenced. Identical colocalization images were also seen when three different anti-CyPA antibodies from diverse suppliers were assayed (goat polyclonal Invitrogen cat #PA5-18463, and two rabbit polyclonals from Invitrogen cat #PA1-025 and GeneTex cat #GTX104698), as well as after co-staining CyPA with the mitochondrial proteins cytochrome c, COX-IV or Tom-20 (data not shown). Fig.1-B shows the colour profiles for CyPA and the mitochondrial marker MitoTracker indicating a high percentage of signal overlapping, and Fig.1-C shows the true colocalization of both signals after subtracting the non-specific background by applying a deconvolution plug-in software.

**Figure 1.**
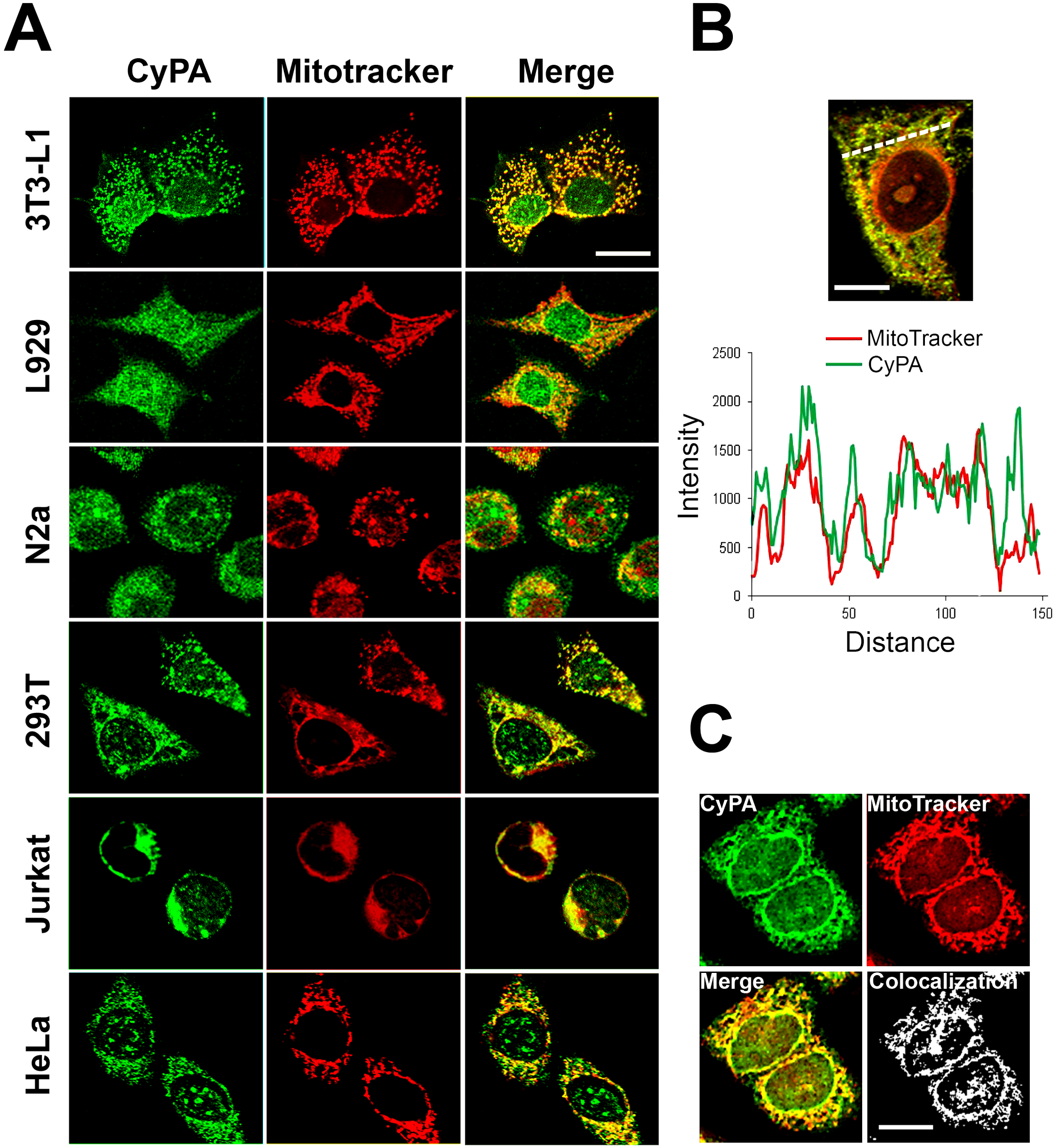
CyPA is a mitochondrial protein. (A) Indirect immunofluorescence images by confocal microscopy show the colocalization of CyPA (green) with the mitochondrial marker MitoTracker® (red) in several cell types. (B) Overlapping of colour profiles for CyPA staining (green) and mitochondria staining (red) in the HeLa cell shown on the top of the plot. (C) Colocalization mask of a z-stack image for 3T3-L1 cells showing the true colocalization of CyPA with mitochondria. Bars= 10 μm.

The amino acid sequence of CyPA does not show a known mitochondrial localization signal that may justify its mitochondrial localization. Therefore, it was analysed the potential requirement of the intrinsic peptidylprolyl isomerase activity of CyPA for this subcellular localization. Fig.2-A shows that the PPIase activity is not required since the inactive point mutant H126Q shows the same colocalization with the mitochondrial probe as the endogenous IMM.

**Figure 2.**
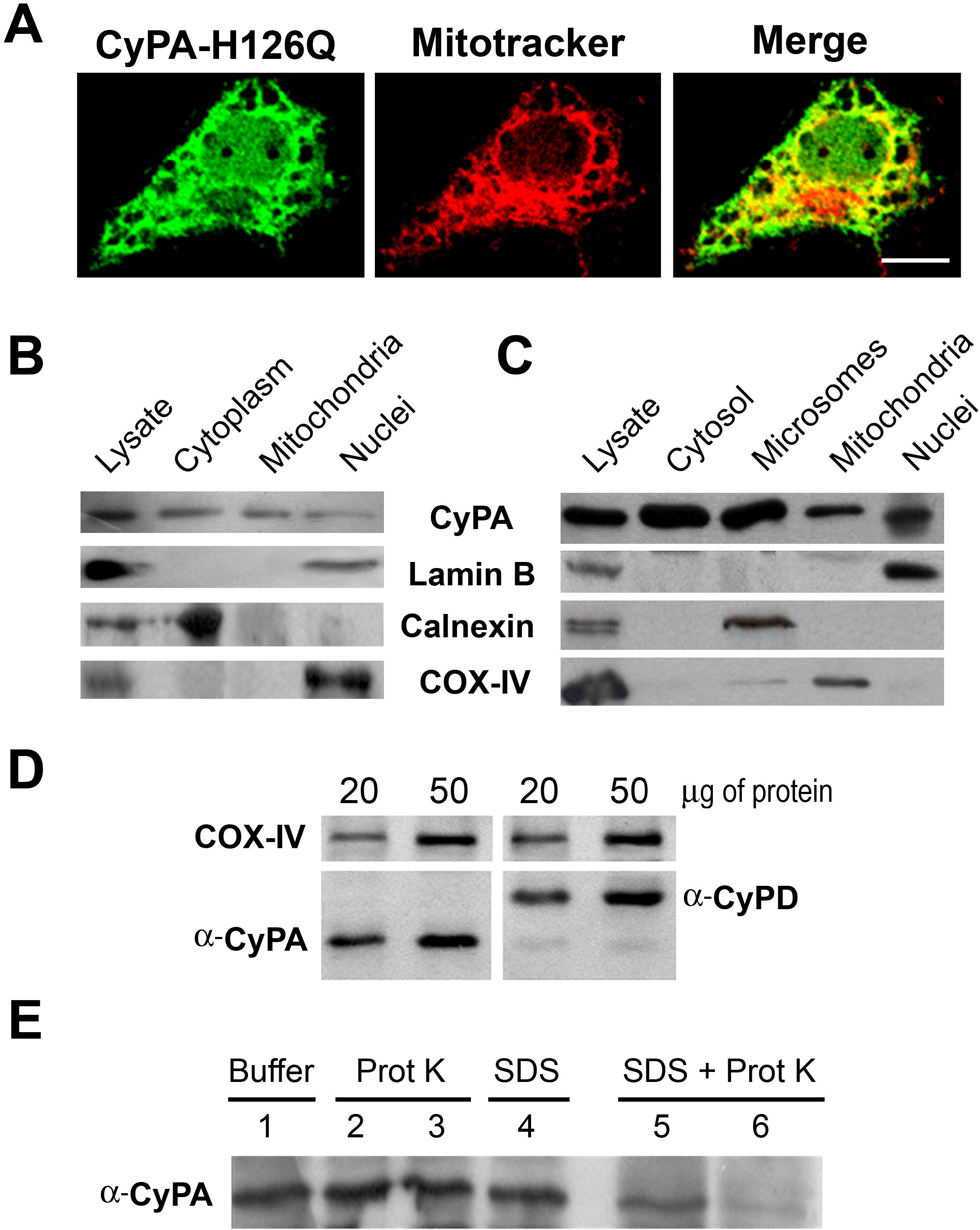
CyPA is intramitochondrial. (A) 3T3-L1 cells were transfected with the PPIase inactive mutant pBEX1-HA-hCyPAH126Q. The subcellular localization of the mutant was visualized with an anti-HA primary antibody followed by a secondary antibody labelled with Alexa-488. Mitochondria were stained with MitoTracker Deep Red-633. (B) Subcellular fractionation of 3T3-L1 cell extracts. (C) Subcellular fractionation of rat liver. (D) 20 μg and 50 μg of total proteins from purified mitochondria of 3T3-L1 cells were resolved in a 15%PAGE/SDS followed by Western blotting for CyPA (left membrane), CyPD (right membrane), and COX-IX as loading control (top membrane). (E) Isolated mitochondria from rat liver were incubated with 1%SDS (lanes 4, 5 and 6), proteinase K (PK) at 5 ng/ml (lanes 2 and 5) or 50 ng/ml (lanes 3 and 6). CyPA was resolved by Western blot.

To confirm the mitochondrial localization of CyPA suggested by confocal microscopy images, mitochondria were purified by biochemical fractionation from both 3T3-L1 cells (Fig. 2-B) and rat liver tissue (Fig.2-C). Lamin B was used as a specific marker of the nuclear fraction, COX-IV as a mitochondrial marker, and calnexin as a microsomal marker in the case of liver extracts, which underwent an additional 100,000x*g* centrifugation to clarify the cytosolic extract due of the high content of low-density lipids (not present in 3T3-L1 extracts). CyPA was detected in all the fractions, including mitochondria. After scanning the protein bands, it was calculated that the mitochondrial CyPA corresponds to approximately 40-45% of the total pool of the IMM present in the extracts of both types of preparations. Inasmuch as CyPD is a mitochondrial IMM that shows high homology with CyPA and regulates the calcium efflux through the permeability transition pore (Mehta, 2018), a control experiment was performed to demonstrate that the anti-CyPA antibody cannot recognize mitochondrial CyPD. Fig.2-D shows Western blots for both proteins from purified mitochondria that have been resolved in a highly resolutive 15%PAGE/SDS. Clearly, the anti-CyPA antibody does not cross-react with CyPD.

To determine whether CyPA is within mitochondria rather than associated to the outer membrane of the organelle, purified mitochondria form 3T3-L1 fibroblasts were treated for 15 min at 4°C with proteinase K (5 ng/ml in lines 2 and 5, or 50 ng/ml in lines 3 and 6). Then, proteins were resolved by Western blot (Fig.2-E). Lanes 2 and 3 show that the presence of CyPA in the mitochondrial pellet remains unchanged after the controlled digestion with the protease. However, the pretreatment of mitochondria with 1% SDS for 10 min degraded the IMM in a protease concentration-dependent manner. This indicates that CyPA is indeed an intramitochondrial factor rather than a protein associated to the outer membrane of the organelle.

### CyPA migrates to the nucleus and shows antiapoptotic properties

Interestingly, CyPA increases its nuclear localization upon the onset of any type of stress. Therefore, it is important to point out that the notorious mitochondrial localization evidenced in fresh cell cultures grown under optimal conditions is lost along the number of passages of the culture when it ages, as well as when even fresh cells become confluent or the medium is not conveniently replaced. These casual observations led us to analyse the subcellular localization of CyPA under several situations of stress (i.e. heat-shock, serum starvation, oxidants, UV light, etc.). As a representative observation among a number of assayed stress situations, Fig. 3-A shows the nuclear translocation of CyPA after the addition to the medium of either aggressor, hydrogen peroxide or staurosporine. Such nuclear translocation shows an average half-life of 2 h for most assayed stimuli, including these two. Therefore, 4 h after the onset of stress, CyPA becomes almost entirely nuclear in all cell types assayed.

**Figure 3.**
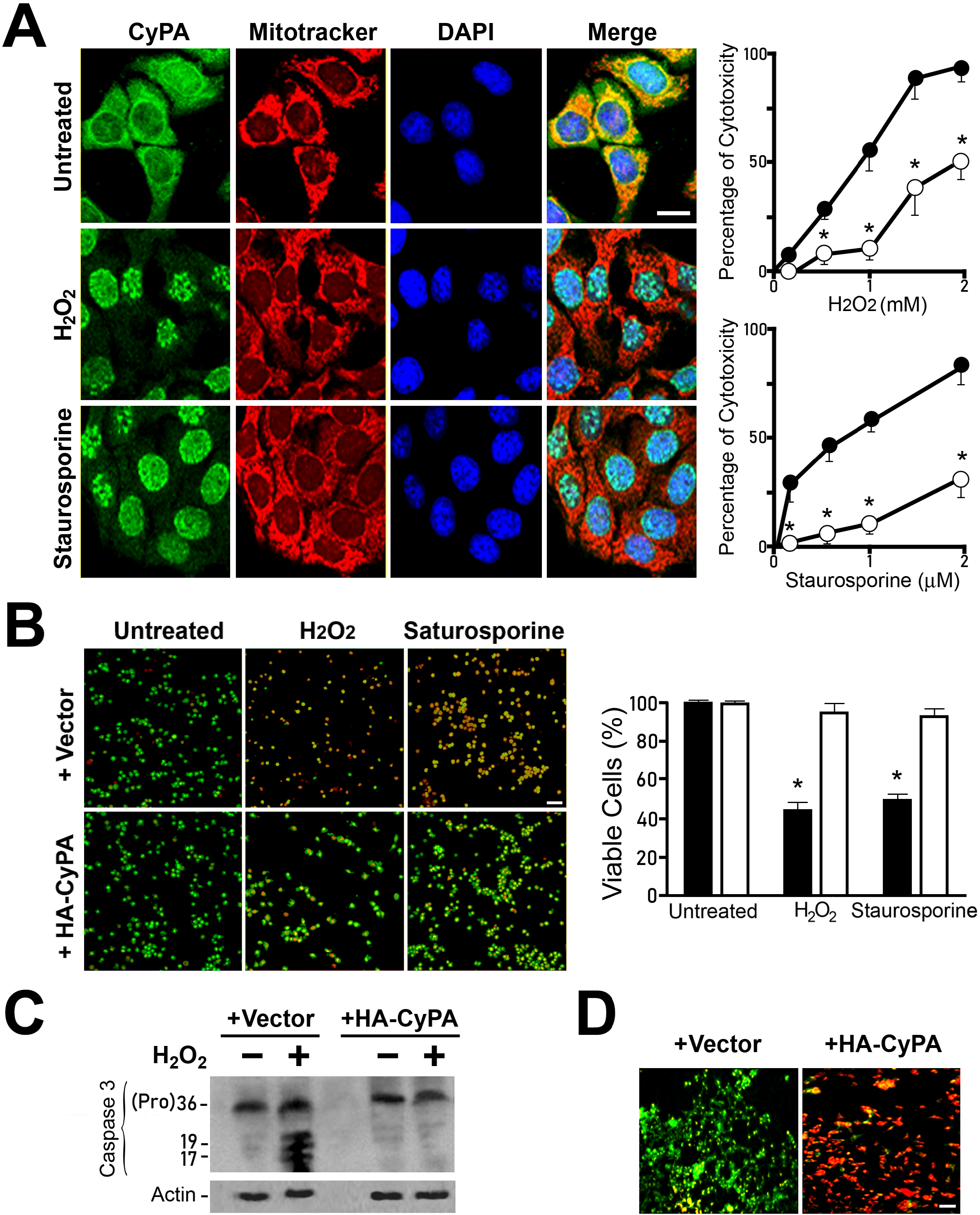
CyPA shows antiapoptotic properties. (A) 293T fibroblasts were treated for 4 h with 100 nM staurosporine or 0.5 mM H_2_O_2_, fixed and stained with MitoTracker (red) or by indirect immunofluorescence for CyPA (green). Images correspond to confocal microscopy (Bar= 10 μm). Plots on the right-side show dose-response curves for the cytotoxicity assay of cells overexpressing HA-CyPA (-●-) versus cells transfected with the empty vector (-○-). *significantly different at p<0.005. The antiapoptotic action of CyPA was also assayed in HA-CyPA 293T overexpressing cells according to: (B) acridine orange/ethidium bromide test after treatement with 100 nM staurosporine; (C) procaspase 3 cleavage upon cell exposure to 0.5 mM H_2_O_2_; and (D) JC-1 cell staining after 10 μM etoposide stimulation. Bars in panels B and D= 100 μm.

We have previously described similar mitochondrial-nuclear translocation for other immunophilin, FKBP51, which also shows antiapoptotic properties associated to such subcellular relocalization. Therefore, the viability of cells overexpressing CyPA or not was analysed after treatment with hydrogen peroxide or staurosporine. The plots shown on the right-side of Fig. 3-A clearly evidence the protective effect of CyPA along the entire range of concentrations of the proapoptotic stimulus. Similar conclusions can be reached after analysing the cells by acridine orange/ethidium bromide test in stauroporine-treated cells (Fig.3-B), the procaspase-3 cleavage after cell exposure to H_2_O_2_ (Fig. 3-C) or cell staining with JC-1, a mitochondrial membrane potential marker, in etoposide-induced apoptosis conditions (Fig. 3-D).

Taken together, all these findings demonstrate that CyPA is a mitochondrial factor that undergoes trafficking towards the nucleus and preserves cell viability when it is overexpressed. It should be noted that CyPA is also a molecular chaperone and its upregulation has been reported under several situations of stress as well as in several types of cancers (melanoma, small cell lung cancer, pancreatic cancer, breast cancer, colorectal cancer, squamous cell carcinoma, etc). A correlation between CypA overexpression and malignant transformation has also been established (Lee, 2010; Nigro et al., 2013).

### CyPA and p23 colocalization

The small acidic cochaperone p23 is usually found associated to the molecular chaperone Hsp90 in most heterocomplexes. Nonetheless, p23 has also Hsp90-independent chaperone activity that can prevent protein aggregation and maintain proteins in a folding-competent state (Freeman et al., 1996). As stated before, molecular chaperones and co-chaperones (including p23 (Madon-Simon et al., 2017; Patwardhan et al., 2013)) are directly involved in the regulation of apoptotic mechanisms in response to accumulation of misfolded proteins, oxidative stress, DNA damage, etc. Because of Fig.3 shows that CyPA undergoes subcellular relocalization upon the onset of stress, comparative indirect immunofluorescence studies were also conducted for CyPA and various molecular chaperones. It was observed that CyPA shows significant colocalization with the Hsp90 cochaperone, p23, especially in neuronal cells. This is an interesting observation since CyPA is not an Hsp90 interactor (Galigniana et al., 2004)[https://www.picard.ch/downloads/Hsp90interactors.pdf]. Fig.4-A shows that p23 undergoes the same subcellular relocalization as CyPA in N2a neuroblastoma cells exposed to H_2_O_2_. The scatter-plot of Fig. 4-B shows a 2D-histogram for both channels where the intensity values of each pixel are plotted against each other. Pixels possessing overlapping signals as it is the case of a true colocalization, plot along a 45° line, i.e. close to the pixels depicted here for the channels of p23 and CyPA (note the excellent linear regression coefficient *r*=0.95).

**Figure 4.**
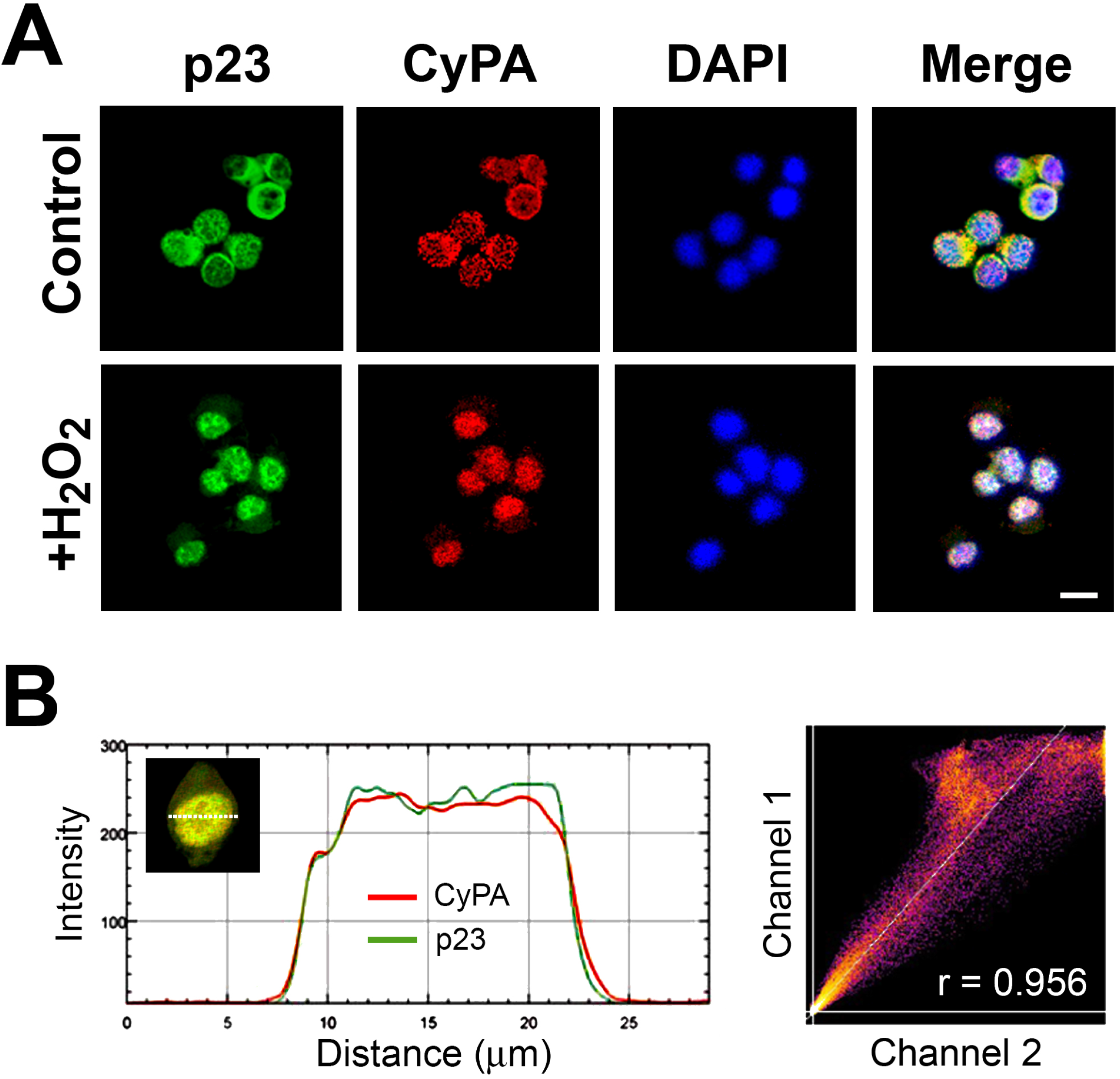
CyPA colocalizes with the cochaperone p23. (A) Indirect immunofluorescence for endogenous CyPA (red) and p23 (green) in N2a neuroblastoma cells. Both proteins comigrate to the nucleus in the presence of 20 μM H_2_O_2_. Bar graph= 10 μm. (B) The cytofluorogram depicts the correlation between the intensities of the two channels of the pixels over distance for the nucleus shown in the insert. The 2D-histogram depicts the intensity values for each voxel when both dyes are plotted against each other in a *z*-stack of the nucleus. The brighter the colour, the more voxels have those two intensity values for their two channels (r= linear regression coefficient).

Because colocalization does not prove the existence of true heterocomplexes, p23 was immunoprecipitated from N2a cell extracts (Fig.5-A). As expected, Hsp90 and Hsp70 were co-immunoprecipitated since p23 belongs to this type of complexes, and CyPA was also pulled-down. To evaluate whether these complexes of p23 and CyPA require the presence of Hsp90, N2a cells were transfected with a hemagglutinin-tagged CyPA, and this exogenously expressed IMM was pulled-down with an anti-HA antibody (Fig.5-B). The co-chaperone p23 was recovered with CyPA in the immunopellet, but free of Hsp90. These results indicate that CyPA and p23 are not associated via Hsp90, and that this chaperone is not require for the assembly of CyPA•p23 complexes. Nevertheless, Hsp90 still forms separate complexes with p23 which are co-immunoprecipitated with an anti-p23 antibody (Fig.5-A), which recognizes the co-chaperone in both types of complexes, Hsp90•p23 and CyPA•p23.

**Figure 5.**
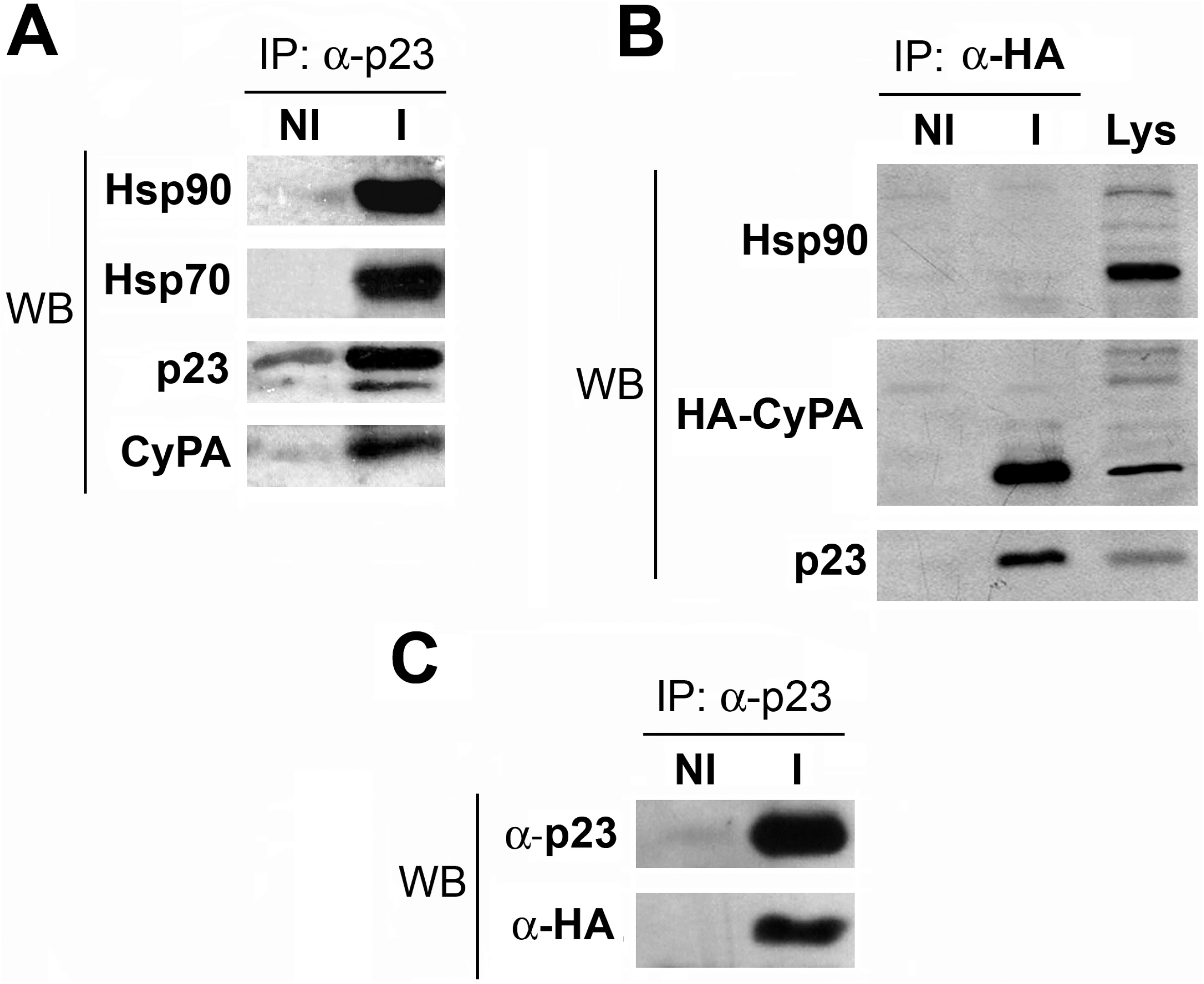
CyPA and p23 interact directly. (A) N2a cytosol was immunoprecipitated (IP) with an anti-p23 IgG (I). A mouse non-immune IgG (NI) was used as control. Proteins were resolved in a 15%PAGE/SDS followed by Western blot (WB) for the indicated proteins. (B) N2a cells transfected with pBEX1-HA-hCyPA were homogenized. The lysate (Lys) was immunoprecipitated (IP) with an anti-HA IgG (I) or a non-immune mouse IgG (NI), and proteins resolved by Western blot as in panel A, except that CyPA was revealed with anti-HA IgG. (C) Cell lysates obtained as in panel B were immunoprecipitated with anti-p23 IgG and Western blotted for the cochaperone and HA.

A piece of straightforward evidence that shows a direct association between CyPA and p23 is the Far Western blot shown in Fig.5-C. Immunopurified p23 was resolved by 15%PAGE/SDS followed by incubation with recombinant GST-CyPA. The resultant complex of the IMM with the co-chaperone was visualized by ECL using an anti-GST antibody and then a secondary counter-antibody tagged to HRP. The development of GST-CyPA on the band of p23 clearly indicates that the interaction of both proteins is direct. It should be remarked that PPIase enzymatic activity of CyPA does not appear to be required for its association with p23 since the blot showed identical results when it was performed in the presence of 1 μM cyclosporine A, a drug that inhibits the PPIase activity (data not shown).

### Overexpression of p23 enhances the antiapoptotic effect of CyPA

With the purpose of investigating the influence of p23 overexpression on the cell response to a proapoptotic stimulus, N2a cells were transfected with the plasmid pRK5MCS-hp23 encoding for human p23 and stimulated with 0.25 μM H_2_O_2_ for 18 h. The percentage of cells showing fluorescence after an incubation with Annexin-V-FITC is shown in Fig.6-A. Clearly, p23 overexpression (see Western blot) exerts a significantly protective action. As expected, such increased level of expression of the cochaperone is reflected in a higher recruitment of p23 to CyPA (see the insert showing co-immunoprecipitations).

**Figure 6.**
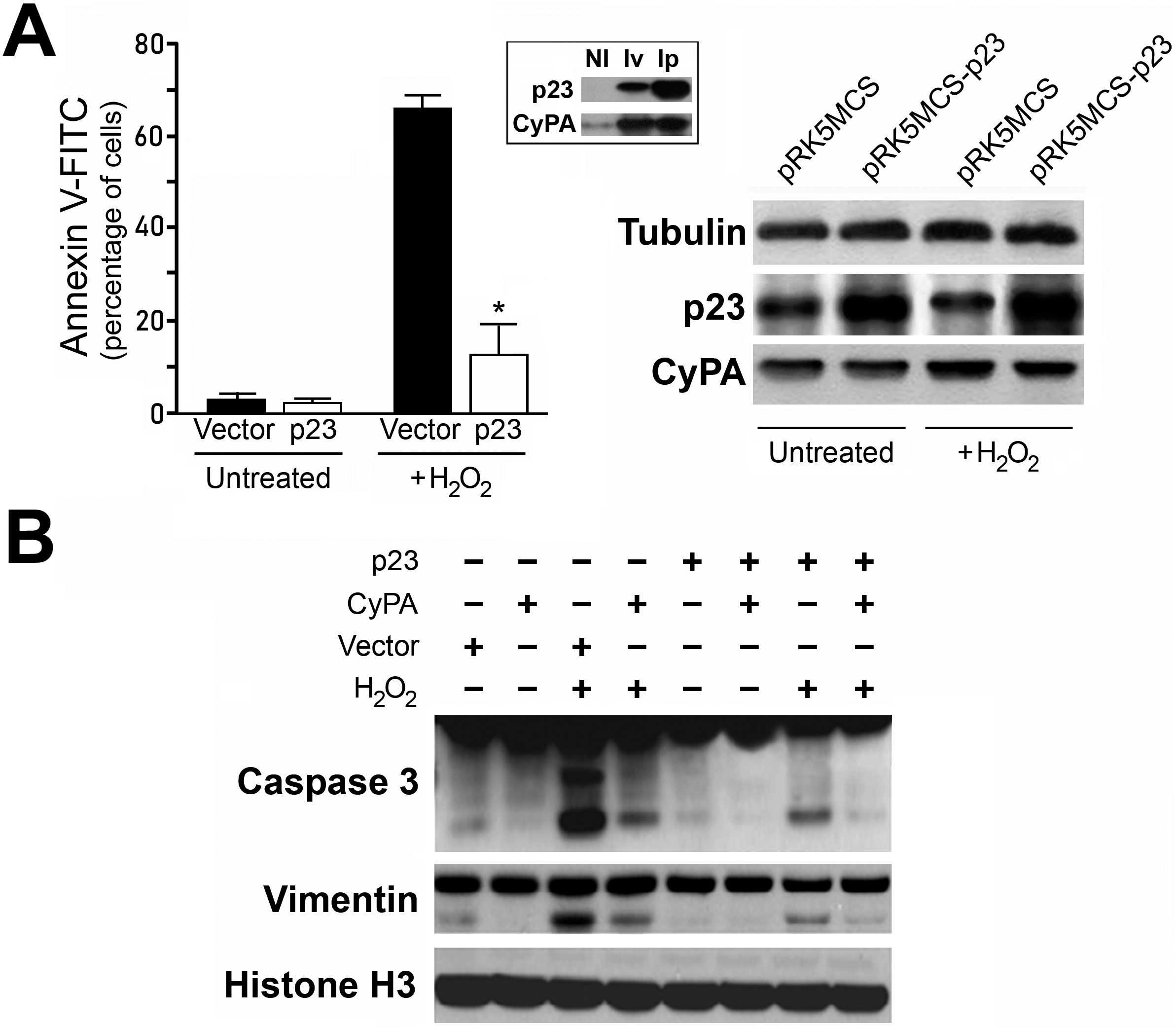
CyPA and p23 overexpression favours anti-apoptotic effect. (A) N2a cells transfected with pRK5MCS-hp23 were exposed to H_2_O_2_ and the percentage of Annexin-V positive cells was quantified by flow cytometry. Results are the mean ± SEM of three independent experiments, each one quantified in triplicate. *Significantly different form cells transfected with empty vector and treated with peroxide at p<0.001. The Western blot on the right-side shows the protein profile for each type of lysate, and the insert shows the greater specific recruitment of p23 in the immunopellet of p23-transfected cells (Ip) versus empty vector transfected cells (Iv). (B) Cells were cotransfected with both proteins, CyPA and p23, and the cleavage of procaspase 3 and the intermediate filament vimentin were evaluated by Western blot after the treatment with H_2_O_2_.

Next, the effect of the overexpression of both CyPA and p23 on the resistance to apoptosis was assayed. Fig. 6-B demonstrates that a high level of expression of both proteins together in cells that were exposed to peroxide greatly attenuates the cleavage of both procaspase 3 and vimentin, whose proteolysis is considered a typical apoptotic response. In other words, the overexpression of both p23 and the IMM, leads to the formation of larger amounts of CyPA•p23 complexes enhancing the protective action of each individual protein in cells exposed to harmful conditions.

## Discussion

In this study it is demonstrated that CyPA is a mitochondrial factor that migrates to the nucleus upon the onset of stress and is capable to exert antiapoptotic actions when overexpressed. It is also shown that CyPA forms complexes with the cochaperone p23, and that high levels of expression of both factors favours the formation of CyPA•p23 complexes enhancing the antiapoptotic effect shown by each individual protein. It is important to note that the stimuli do not generate cell death by autophagy or necrosis, but apoptosis as it is shown by the different assays accepted to demonstrate this mechanism of cell death (i.e. procaspase and vimentin cleavage, annexin-V binding to plasma membrane, changes in the mitochondrial potential evidenced with JC-1, etc.). Remarkably, the subcellular localization of CyPA as well as its nuclear translocation triggered by stress was a phenomenon evidenced in all cell types assayed and using both confocal microscopy studies and standard biochemical fractionations, suggesting it is not restricted to a specific cell type. Furthermore, the mitochondrial localization of CyPA was also evidenced in rat liver mitochondria (Fig.2) and other organs such as brain (data not shown).

Interestingly, the novel association of CyPA with the small acidic cochaperone p23 does not require Hsp90, a chaperone that itself usually requires p23 to stabilize its heterocomplexes (Forafonov et al., 2008; Freeman et al., 1996; Pratt et al., 2004b). Importantly, p23 has also been indirectly associated with antiapoptotic actions in other studies. An exacerbated instability of p23 leads to breast and cervical carcinoma cells to apoptosis-mediated death (Patwardhan et al., 2013), suggesting a protective role for p23. Accordingly, cancer and metastatic cells show higher levels of p23 expression (Mollerup et al., 2003; Oxelmark et al., 2006), the prognosis is worse in prostate cancer patients with increased nuclear localization of p23 (Cano et al., 2015), and its overexpression protect cells against the deleterious actions of Hsp90 inhibitors (Forafonov et al., 2008). A similar situation is caused by CyPA upregulation in malignant transformation and several types of cancers [1, 29]. Therefore, the potentiation of CyPA•p23 as a novel antiapoptotic complex may be linked to improved survival skills of cancer cell, such that the disruption of this association by drugs deserves to be attempted as a novel putative target against cancer.

In addition to its role in protein folding, CyPA has also been involved in intracellular trafficking (Galigniana et al., 2004; Zhu et al., 2007), signal transduction (Sun et al., 2014), and transcriptional regulation (Xie et al., 2019). Other cyclophilins were cloned and characterized after CyPA identification, among them CyPD (also known as PPIase-D), which shows 75% structural homology with CyPA (Mehta, 2018) and is an IMM linked to the inner mitochondrial membrane in the mitochondrial permeability transition pore. In contrast to CyPA oligomeric properties, CyPD is an Hsp90-associated partner and requires this chaperone (Kang et al., 2007). Although it binds cyclosporine A, its biological functions are independent of the PPIase activity (Scorrano et al., 1997). In contrast to the observed effects of CyPA overexpression, increased levels of CyPD like those shown in most neurodegenerative diseases leads to cell death due to colloidal osmotic swelling of the mitochondrial matrix, dissipation of the inner membrane potential, generation of ROS, and the release of many proapoptotic proteins and procaspases (Zhang et al., 2015). Importantly, as opposite to mitochondrial CyPD that is a protein associated to membranes with a role in the regulation of a calcium pore, mitochondrial CyPA is not confined to the organelle and can fully accumulate in the nucleus. In other words, our observations suggest that regardless of their high homology, CyPA and CyPD do not seem to have redundant functions.

Like most members of the subfamily, CyPA is also a stress-inducible protein. Therefore, it is likely that its protective role in the stress response could be an extension of its intrinsic molecular chaperoning nature under normal conditions, a property that is especially exacerbated during harmful circumstances. This would favour associations of CyPA with key members of the chaperone family, for example, the cochaperone p23. Moreover, the beneficial association of CyPA and p23 may be a cardinal requirement for their involvement in the structural transformation that ultimately leads to mitochondria stabilization and the antiapoptotic effect.

## Materials and Methods

### Reagents and plasmids

Staurosporine, proteinase K, MTT, p-formaldehyde, and hydrogen peroxide (cat #S180505-R22-4) were from Sigma (St. Louis, MO, USA). Acetone was from Merck (Darmstadt, Germany). Vectashield^®^ antifade mounting medium for microscopy was from Vector Laboratories Ltd. (Peterborough, UK). All cell types used in this study were purchased from ATCC (Manassas, VA, USA). DMEM and OPTI-MEM culture media were from Invitrogen (Waltham, MA, USA).All plasmids encoding for human CyPA (pGEX-2T-GST-CyPA, pBEX1-HA-hCyPA, and the inactive mutant pBEX1-HA-hCyPAH126Q) were a generous gift from Dr. Philippe A. Gallay (Scripps Research Institute, La Jolla, CA, USA). The plasmid pRK5MCS-hp23 encoding for human p23 was kindly provided by Dr. Theo Rein (Max Planck Institut für Psychiatrie, Munich, Germany).

### Antibodies

The AC88 mouse monoclonal IgG against Hsp90 and the N27F3-4 anti-72/73-kDa heat-shock protein monoclonal IgG (anti-Hsp70) were from StressGen (Ann Arbor, MI). The JJ3 mouse monoclonal IgG against p23 was from Affinity BioReagents (Golden, CO). The 6E2 mouse IgG anti-hemagglutinin tag and rabbit IgG anti-caspase 3 were from Cell Signaling (Danvers, MA, USA). The anti-CyPA goat polyclonal (cat #PA5-18463) and commercial rabbit antiserum (cat #PA1-025) were purchased from Invitrogen (Waltham, MA, USA). The rabbit polyclonal IgG anti-CyPA (cat #GTX104698) was from GeneTex (Irvine, CA, USA). Rabbit antiserum anti-CyPD was a kind gift from Dr. Yolanda Sopena (University of Buenos Aires). The rabbit polyclonal antiserum against CyPA was prepared in the Scripps Laboratory of Dr. Philippe Gallay by immunization with recombinant human CyPA protein (Saphire et al., 1999). Anti-GST rabbit antiserum was a generous gift from Dr. Martín Monte (University of Buenos Aires). Antibodies against cytoskeletal proteins (vimentin, actin and β-tubulin were from Sigma (St.Louis, MO, USA). The mitochondrial marker MitoTracker^®^, JC-1 dye, annexin-FITC, and all fluorescent secondary antibodies were from Molecular Probes (Eugene, OR, USA). Rabbit monoclonal IgG anti-histone H3 was from Upstate (Syracuse, NY, USA). Goat polyclonal IgG anti-lamin B, mouse monoclonal IgG against cytochrome c, rabbit polyclonal IgG anti-Tom-20, and goat polyclonal IgG anti-lamin B, and rabbit polyclonal IgG anti-calnexin were from Santa Cruz Biotechnology (Dallas, TX, USA). Anti-COX-IV was purchased from Abcam (Cambridge, UK). Immunoprecipitation assays were performed as described in previous publications (Mazaira et al., 2020; Piwien-Pilipuk et al., 2002).

### Microscopy studies

Cells were grown on coverslips and processed for indirect immunofluorescence as described in previous studies (Presman et al., 2006; Salatino et al., 2006; Toneatto et al., 2013). Briefly, the cells were fixed at room temperature for 1 h in a freshly prepared solution of 4% p-formaldehyde in PBS, permeabilized for 5 min in cold (−20°C) acetone. After washing the coverslips with PBS buffer, they were preincubated for 1 h at room temperature in high ionic strength solution (HISS) buffer (20 mM Tris at pH 8.0, 0.63 M NaCl, 0.1% Tween 20, and 3.5% bovine serum albumin) supplemented with 1% horse serum, and then incubated for 2 h at 4 °C with 1/250 dilution (in HISS buffer) of primary antibody followed by 1 h incubation at room temperature with 1/200 dilution of secondary antibody. The coverslips were mounted in a glycerol based medium with an antifade solution. Confocal microscopy images were acquired with a Nikon Eclipse-E800 confocal microscope using a Nikon DSU1 camera with ACT-2U software. Co-localization analyses were performed using the co-localization plug-in of the Fiji program (v.1.45) (NIH; Bethesda, MA, USA), which uses a range of algorithms such as co-localization thresholds, Pearson’s linear correlation coefficient, overlap and Manders’ coefficients (Manders et al., 1993). We collected confocal z-series of cells (40 optical slices, airy unit ¼ 1 airydisc at 0.25 mm intervals using 63x objective), and then images were deconvolved using Huygens compute engine 3.5.2p3 64b (closed platform). Finally, images were imported to the Fiji program to determine the degree of overlapping.

### Cell fractionation and proteinase K digestion

Mitochondria were isolated from rat liver or cell extracts following a standard procedure for biochemical fractionation (Gallo et al., 2011). In order to determine whether CyPA is intramitochondrial, isolated mitochondria were permeabilized by pretreatement for 10 min at 4°C with 1% SDS, and then incubated with 5 or 50 ng/ml proteinase K for 15 min (Kang et al., 2007). The presence of CyPA was resolved by Western blot in a 15%PAGE/SDS.

### Apoptosis and cell viability tests

Procaspase and vimentin cleavage was assayed as previously described (Gallo et al., 2011). Acridine orange/ethidium bromide test was achieved according to Liu et al. (Liu et al., 2015). Cell viability was assayed as previously reported (Colo et al., 2008; Lagadari et al., 2016). Etoposide-induced apoptosis in human 293T fibroblasts was induced according to Attardi et al. (Attardi et al., 2004). Cell staining with JC-1, annexin-FITC and MitoTracker markers was accomplished as previously described (Gallo et al., 2011).

### Far Western Blot

Proteins from 3T3-L1 fibroblast lysates extracted with Cytobuster protein extraction reagent (Novagen^®^, Merck). After cell disruption for 10 min at room temperature, the extracts were centrifuged at 10,000x*g* for 10 min. The co-chaperone p23 was immunoprecipitated with JJ3 anti-p23 mouse monoclonal IgG, washed three times with TEGMo buffer (10 mM TES at pH 7.6, 50 mM NaCl, 4 mM EDTA, 10% v/v glycerol, and 20 mM Na_2_MoO_4_), and proteins were resolve by a 15% PAGE/SDS, transferred to Immobilon-P membrane, and incubated with a recombinant GST-CyPA solution obtained from IPTG-induced *Escherichia coli* as described in previous studies (Galigniana et al., 2004; Galigniana et al., 2001). After washing the membrane, the attachment of GST-CyPA to p23 was visualized by ECL with anti-GST antiserum/donkey anti-rabbit IgG-HRP.

## Acknowledgements

This study was possible thanks to the economical supported by grants from the Universidad de Buenos Aires (UBACYT 20020170100558BA) and the Agencia Nacional de Promoción Científica y Tecnológica (PICT 2016-0545 and PICT 2018-0546). The authors would like to express their gratitude to Dr. Philippe Gallay from the Scripps Research Institute (La Jolla, CA, USA) and Dr. Theo Rein from the Max Planck Institut für Psychiatrie (Munich, Germany) for the generous provision of the plasmids used in this work.

